# Differentiable Partition Function Calculation for RNA

**DOI:** 10.1101/2023.01.30.526001

**Authors:** Marco Matthies, Ryan Krueger, Andrew Torda, Max Ward

**Author notes:** These authors contributed equally to this work (Marco Matthies, Ryan Krueger).

## Abstract

Ribonucleic acid (RNA) is an essential molecule in a wide range of biological functions. In 1990, McCaskill introduced a dynamic programming algorithm for computing the partition function of an RNA sequence. This forward model is widely used for understanding the thermodynamic properties of a given RNA. In this work, we introduce a generalization of McCaskill’s algorithm that is well-defined over continuous inputs and is differentiable. This allows us to tackle the inverse folding problem—designing a sequence with desired equilibrium thermodynamic properties—directly using gradient optimization. This has applications to creating RNA-based drugs such as mRNA vaccines. Furthermore, it allows McCaskill’s foundational algorithm to be incorporated into machine learning pipelines directly since we have made it end-to-end differentiable. This work highlights how principles from differentiable programming can be translated to existing physical models to develop powerful tools for machine learning. We provide a concrete example by implementing an effective and interpretable RNA design algorithm.

## 1 Introduction

Ribonucleic acid (RNA) is fundamentally important to biology [1] and carries out many of the processes of life alongside DNA and protein such as transcription and translation [2], catalyzing reactions [3], gene regulation [4], and much more. Many of these functions require specific structures [5], so effort has gone into computational methods for predicting structures [6], which are now widely used [7, 8]. These are based on the nearest neighbor model [9, 10, 11, 12], which was found empirically through numerous experiments and predicts the thermodynamics of RNA secondary structures. A fundamental part of these methods is the computation of the *partition function*.

There exists a dynamic programming algorithm for computing the partition function of an RNA sequence under the nearest neighbor model [13]. This algorithm is widely used for computing various statistical properties of an RNA in thermodynamic equilibrium such as individual base-pair probabilities, the probability of folding into a given secondary structure, and heat capacity [14].

This approach is in stark contrast to recently developed deep learning methods for studying proteins (e.g., AlphaFold) [15, 16]. First, deep learning models of protein structure and function do not typically operate directly on thermodynamic quantities or explicitly predict ensemble statistics. Second, unlike deep learning models, the calculation of the partition function of an RNA sequence is only well-defined for a discrete sequence and therefore is not differentiable. This limits downstream applications of the model. For example, end-to-end differentiable models of protein structure enable new methods for inverse folding. The inverse folding problem (often also called the design problem) inverts the structure prediction problem. Instead of predicting the structure given the sequence, we must find a sequence that folds into a target structure. A differentiable folding algorithm can be inverted by starting from any sequence and optimizing the sequence using gradients propagated through the forward model [17, 18]. An end-to-end differentiable RNA partition function would enable this method for inverse RNA folding.

An end-to-end differentiable RNA partition function is important for other tasks too. It extends a classical and widely used algorithm, which can now be incorporated into machine learning pipelines, for example as a loss function in deep learning. In addition, it can be used to optimize the parameters of the nearest neighbor model in a similar way to inverse folding, which is an important but challenging task [19].

In this work, we adapt the classical dynamic programming algorithm for computing an RNA’s partition function by making it end-to-end differentiable. To achieve this, the algorithm must (i) operate on a continuous input and (ii) comprise of differentiable operations. To satisfy (i), the input to our algorithm is a *probabilistic* sequence (i.e. a probability distribution of sequences) and we introduce new thermodynamic quantities for describing the partition function of such a sequence. Crucially, we then modify the existing algorithm to compute this partition function by calculating intermediate Boltzmann-weighted sums that are conditioned on the base identities of the subsequence in question. We satisfy (ii) by implementing our method in JAX [20].

To demonstrate the effectiveness of our method, we use it for inverse folding. The partition function can be used to calculate the probability of a structure. The probability of a structure is known to be a good guide for inverse folding [21]. Gradients with respect to the sequence distribution can be computed, which lets us optimize the probability by gradient descent. Success in increasing the probability indicates that our method is correct, and that the gradients we calculate are meaningful.

## 2 Methods

In this section, we present the definition of a partition function for a probability distribution of sequences and our algorithm for computing this quantity in a differentiable fashion. We then present a small set of inverse folding experiments performed to test our approach.

### 2.1 Partition Functions for Probabilistic Sequences

The partition function of a system is defined as

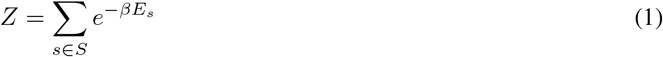

where *S* is the set of all microstates, *β* is the inverse temperature, and *E*_*s*_ is the total energy of the system in microstate *s*.

In the context of RNA folding, the set of possible microstates *S* for a given sequence is defined as all possible secondary structures under the model. A secondary structure is typically described as a properly nested set of parentheses where indexes are permitted to be unpaired [6, 13]. Formally, for a sequence of length *n*, a secondary structure is a set of pairs (*i, j*) ∈ {1 … *n*} ^2^ such that *i < j* and there are no two pairs (*i, j*) and (*k, l*) where *i < k < j < l* and there are no pairs with shared indexes.

For a fixed RNA sequence, the partition function described by Equation 1 corresponds to the sum of Boltzmann weights over all secondary structures. We refer to this fixed-sequence, variable-structure partition function as the *structure partition function*, denoted *Z*_struct_. For a sequence *q*, the structure partition function is defined as

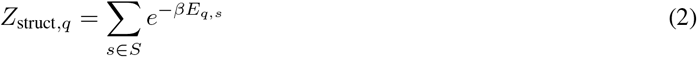

where *E*_*q,s*_ denotes the energy of sequence *q* with structure *s* under the nearest neighbor model. This is what McCaskill’s algorithm computes [13].

In this work, we introduce two new thermodynamic quantities: the *sequence partition function* and the *structure-sequence partition function* denoted *Z*_seq_ and *Z*_ss_, respectively. Consider a probability distribution of sequences of length *n*,

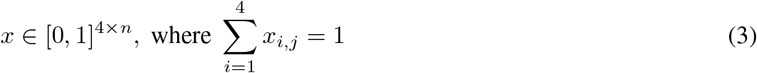

where the value at each of the four indices at a given position *j* corresponds to the probability of the the nucleotide at that position having identity A, C, G, or U. For a given discrete sequence *q* ∈ *Q* where *Q* is the set of all 4^*n*^ sequences of length *n*, the probability of sampling *q* from *x* is the product of all corresponding base identities at each position. Intuitively, this can be thought of as a probabilistic, or “fuzzy” sequence. Crucially, a probabilistic sequence is continuous and can be analysed using a gradient.

For a given secondary structure *s*, we define the sequence partition function as the weighted sum of Boltzmann weights over all sequences *q* ∈ *Q* assuming structure *s*:

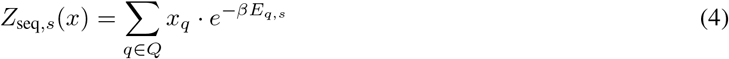

where *x*_*q*_ is the probability of sampling sequence *q* from *x*. Similarly, we define the structure-sequence partition function:

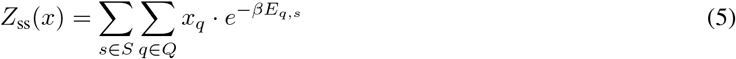

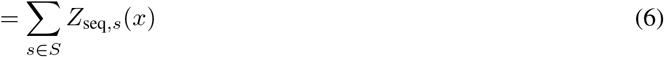

See Section 2.3 for an example of how this partition function can be interpreted in the context of thermodynamic properties.

### 2.2 Calculation of Structure-Sequence Partition Function

We modify McCaskill’s algorithm [13] so that it is differentiable. This involves two major conceptual changes. First, the input must be made continuous, and is therefore described by a probability distribution (see Equation 3). We describe new recursions to work with a probability distribution. Second, all the operations need to be differentiable. We re-implemented the energy functions to be differentiable, and the recursions to contain only differentiable operations. This was achieved using JAX [20].

We implemented the nearest neighbor energy model with three main energy functions [9]. The model gives each loop a free energy contribution and the sum of contributions is the free energy of the structure. A loop is a region enclosed by base pairs and each kind of loop has a different energy function. See Figure 1 for an example. We modify the energy functions to work with explicit base identities to support a probabilistic sequence.

**Figure 1:**
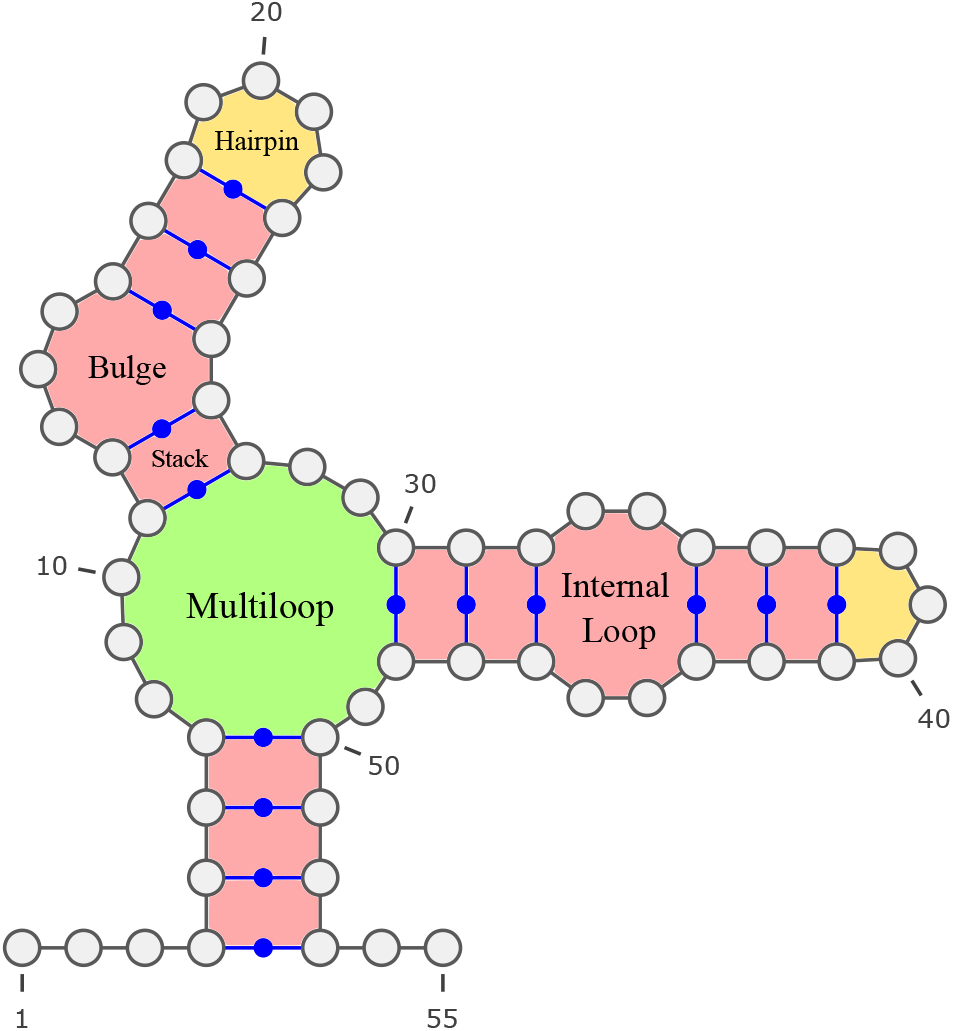
An example RNA structure showing the nearest neighbor model. One-loops are depicted in yellow, two-loops in red, and multi-loops in green. Examples of specific nearest neighbor model motifs are given. This image was generated with the help of VARNA [22].

The one-loop(*b*_*i*_, *b*_*j*_, *i, j*) function gives the free energy contribution of a loop closed by a single pair *i, j*, also called a hairpin loop. The parameters *b*_*i*_ and *b*_*j*_ refer to the bases (A, C, G, or U) at positions *i* and *j*.

Next, we have the two-loop(*b*_*i*_, *b*_*j*_, *b*_*k*_, *b*_*l*_, *i, j, k, l*) function, which gives the free energy of a loop formed by the two pairs *i, j* and *k, l*. This captures stacks, bulge, and internal loops in the nearest neighbor model. The parameters *b*_*i*_, *b*_*j*_, *b*_*k*_, *b*_*l*_ describe the base identities of the corresponding locations.

Finally, the model uses an affine approximation of loops closed by 3 or more pairs. These are called multiloops. A multiloop closed by *P* pairs and containing *U* unpaired nucleotides is assigned a free energy contribution of *M*_*i*_ + *M*_*p*_ · *P* + *M*_*u*_ · *U*.

We have left out several complexities in the energy model for brevity. These include terminal mismatches, dangling ends, helix end penalties, special hairpins, and special internal loops. These amount to tedious case-work and the core ideas of the algorithm are clearer without them. Details can be found in our code.

For convenience, we define a Boltzmann function,

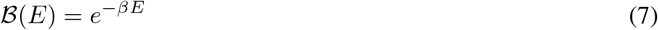

which scales a given energy term *E*.

The structure-sequence partition function is computed by dynamic programming described by three functions. The first computes the external loop. Define ℰ (*i*) as the partition function of the fragment of nucleotide [*i, n*) assuming the nucleotides are indexed from 0 to *n* − 1. The base case is ℰ(*n*) = 1. Define 𝒫(*b*_*i*_, *b*_*j*_, *i, j*) as the partition function of the fragment [*i, j*] conditioned on *i* being base *b*_*i*_ and *j* being base *b*_*j*_ and assuming *i, j* form a pair. The base cases are 𝒫(*b*_*i*_, *b*_*j*_, *i, j*) = 0 when *i* ≥ *j*. Define ℳ(*p, i, j*) as the partition function of the fragment [*i, j*] assuming it is in a multiloop, and the fragment contains at least *p* pairs closing the multiloop. The base cases are ℳ(0, *i, j*) = 1 when *i > j* and ℳ(*p >* 0, *i, j*) = 0 also when *i > j*.

We can compute ℰ by summing over the cases where *i* is not paired, and where *i* is paired.

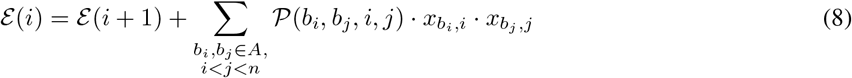

Recall that *x* is the probabilistic sequence and 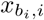 denotes the probability that the sequence is base *b*_*i*_ at location *i*. These probabilities must be multiplied through. Note that there are four combinations each for *b*_*i*_ and *b*_*j*_ that we sum over corresponding to the alphabet of nucleotides *A* = {A, C, G, and U}. In combination, we also sum over all valid right endpoints *j* for a pair *i, j*.

Observe that the partition function itself is defined by ℰ(0).

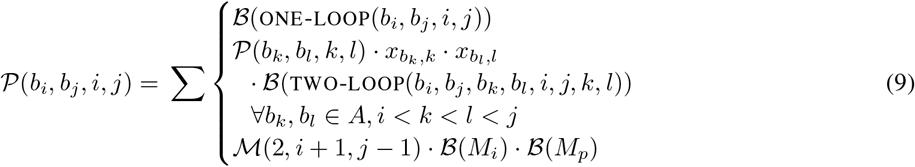

The recursions for 𝒫 sum over three types of case. The first is the one-loop closed by just the pair *i, j*. The second is two-loops closed by the two pairs, *i, j* and some *k, l* that is fully contained inside *i, j*. Note that in the second case we must consider all options for *k, l* and the bases *b*_*k*_, *b*_*l*_ at those locations. Furthermore, that we only multiply by the probabilities 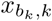 and 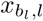 since 𝒫(*b*_*i*_, *b*_*j*_, *i, j*) is conditional on *b*_*i*_ and *b*_*j*_. The third and final case is for multiloops closed by *i, j*. Observe that at least 2 additional pairs are required to make a multiloop, which is why ℳ (2, *i* + 1, *j* − 1) is used.

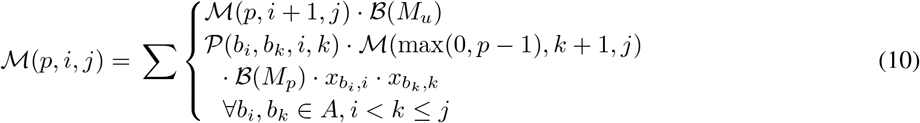

The recursions for ℳ sum over two cases similar to those for ℰ. The first is when position *i* is not paired. The second is when *i* is in a pair *i, k*. In the second case, we must sum over all options for *b*_*i*_, *b*_*k*_ and *k*.

Our implementation targets parity with ViennaRNA 2.5.0 [7]. ViennaRNA is the most widely used RNA folding software pacakge. By default, ViennaRNA double counts dangling ends and terminal mismatches to approximate coaxial stacking. We leave out a detailed discussion of how we implemented this, since it involves subtle casework. However, it is worth pointing out that this significantly complicates the recursions for multiloops, ℳ, and leads to high memory usage. Ironically, the behaviour of ViennaRNA is an approximation designed to make their recursions simpler, but when they are redesigned to the structure-sequence partition function, it makes them more complicated. We expect that the memory usage and speed will improve under a more correct (non-approximate) folding model.

The complexity of our algorithm is the same as McCaskill’s [13] requiring *O*(*n*^4^) time. It could be optimized to *O*(*n*^3^) using speedup by Lyngsø [23]. Also, JAX allowed us to run our algorithm in parallel on the GPU. Given sufficient cores, our algorithm runs in *O*(*n*) time, since all table values for a given row indexed by *i* can be computed in parallel so long as the table values for rows with larger indexes have already been computed, and there are only *n* rows in total.

### 2.3 Test Case: Inverse Folding

A classical result of statistical mechanics is that the probability of a microstate *s* ∈ *S*, where *S* is the set of all possible microstates of a system, is proportional to its Boltzmann weight,

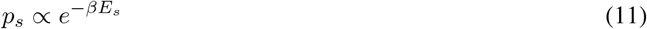

where *β* is the inverse temperature and *E*_*s*_ is the energy of *s*. Since the total probability of all microstates is equal to 1,

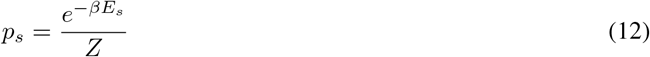

So, the probability of a discrete RNA sequence *q* folding into some structure *s* ∈ *S* is

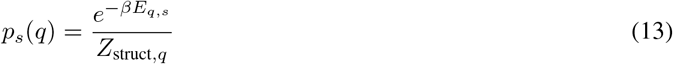

Given the thermodynamic quantities introduced in Section 2.1, we can define an analog of Equation 13 for a probabilistic sequence *x*:

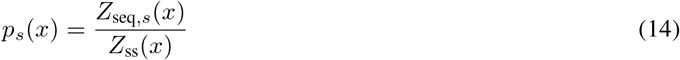

Intuitively, this can be interpreted as the probability that a sequence sampled from the distribution *x* will fold into secondary structure *s*. Note that in the case that *x* is a one-hot encoded vector, Equation 14 reduces to Equation 13.

To validate our method, we perform a small set of inverse folding experiments on structures from the Eterna100 [24], a well known benchmark for inverse RNA folding. We consider all structures from the Eterna100 of length at most 50 given computational limitations of our implementation. For each structure *s*, we apply simple gradient descent over the probabilistic sequence *x* to maximize *p*_*s*_(*x*). In practice, we define our loss function as ℒ_*s*_(*x*) = − log(*p*_*s*_(*x*)). To ensure that the normalization condition in Equation 3 is satisfied for each gradient update, we apply the softmax operator to the sequence logits at each step. All probabilistic sequences were initialized with uniformly distributed base identities at each position. Each design experiment was conducted using 200 iterations of gradient descent with the RMSProp optimizer [25] and a learning rate of 0.1.

The result of this optimization is, by construction, a distribution over sequences and not a single discrete sequence. In our examples, the softmax of sequence logits converged to a one-hot encoded sequence. Though not used in this work, convergence to a discrete sequence can be enforced via the method developed by Roney and Ovchinnikov for inverse protein design with AlphaFold [17].

## 3 Results

We sought to validate the utility of the gradients calculated from the procedure described in Section 2.2 for numerical optimization. To do so, we conducted inverse folding experiments per the method described in Section 2.3 on all 18 structures of length at most 50 from the Eterna100. The results of all 18 optimizations are presented in Table 1. The post-compilation runtime of a single gradient step ranged from 0.06 − 2.67 seconds on an 80 GB NVIDIA A100 GPU.

**Table 1:**
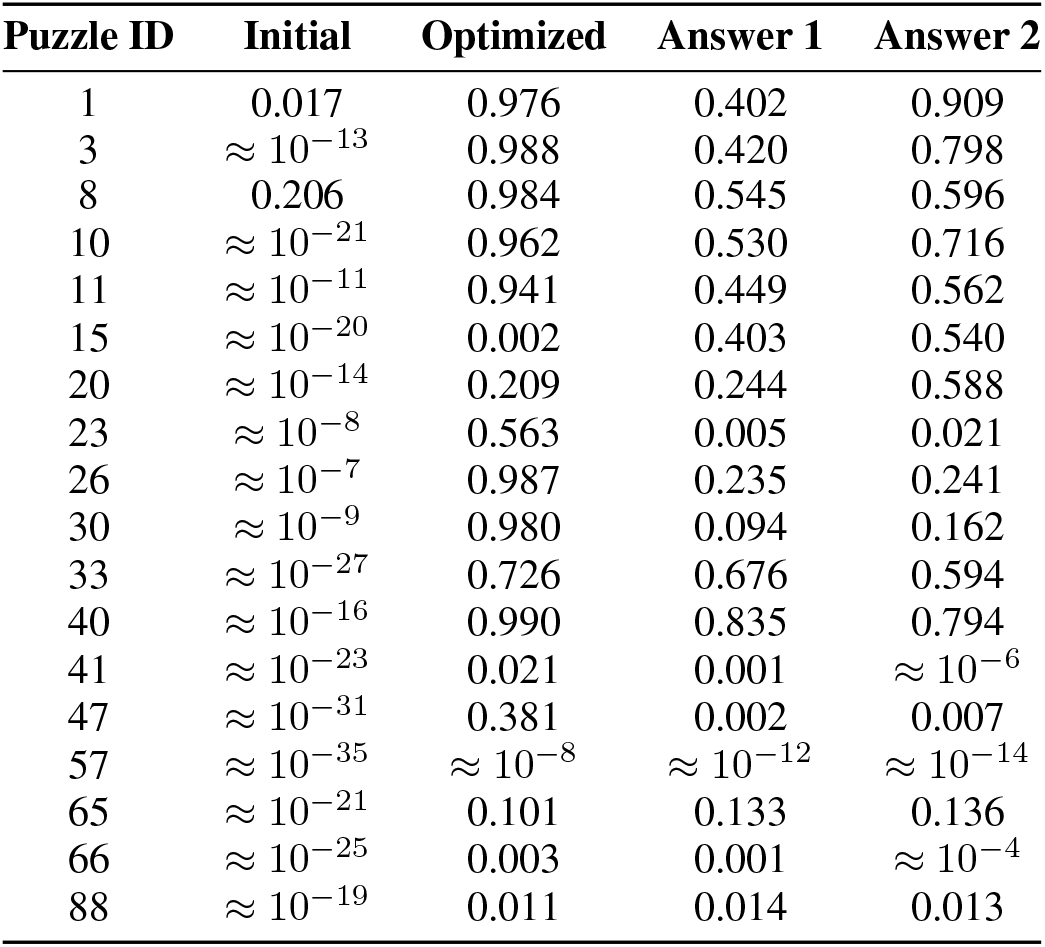
The results of the inverse folding procedure described in Section 2.3 on all Eterna100 structures of length at most 50. Each structure is identified by its ID in the Eterna100 dataset. The “Initial” column corresponds to the probability that a sequence sampled from the initial sequence distribution will fold into the desired structure. The “Optimized” column corresponds to the same probability for the optimized sequence distribution after 200 gradient updates. “Answer 1” and “Answer 2” correspond to the probabilities of the two Eterna100 reference answers folding into the target structure.

In all cases, the initial probability distribution of sequences was improved by gradient descent, as measured by the probability of sampling a sequence that folds into the target structure. In most cases, the probability improved by many orders of magnitude. In 14 of the 18 cases, the final probability distributions (all of which converged to a discrete sequence) outperform both reference answers provided in the Eterna100 dataset. In the remaining four cases (i.e., puzzles 15, 20, 65, and 88), visual inspection of the final probability distributions suggests that the optimization procedure arrived at local minima. Additionally, two of these four cases (i.e. puzzles 57 and 88) are known to be unsolvable using the Vienna 2 parameters [26].

In all of our experiments, normalized sequence entropy drops quickly over the first 30 steps and sequence distributions settle down to a preferred base at each position. In many cases though we see subsequent smaller changes to base identities and corresponding improvements to target probability after many steps of optimization and even after phases of seemingly little change. In Figure 2 we visualize the successful optimization run for puzzle 10 (“Frog Foot”). After 120 optimization steps the U14-A19 positions, which are base-paired in the target structure, change to a C-G base pair via a U-G intermediate, resulting in an an additional small improvement to target probability. Other changes at positions unpaired in the target structure are the intermittent A to U switch at position 6 during steps 25–175 and the U to A change at position 10 around step 175.

**Figure 2:**
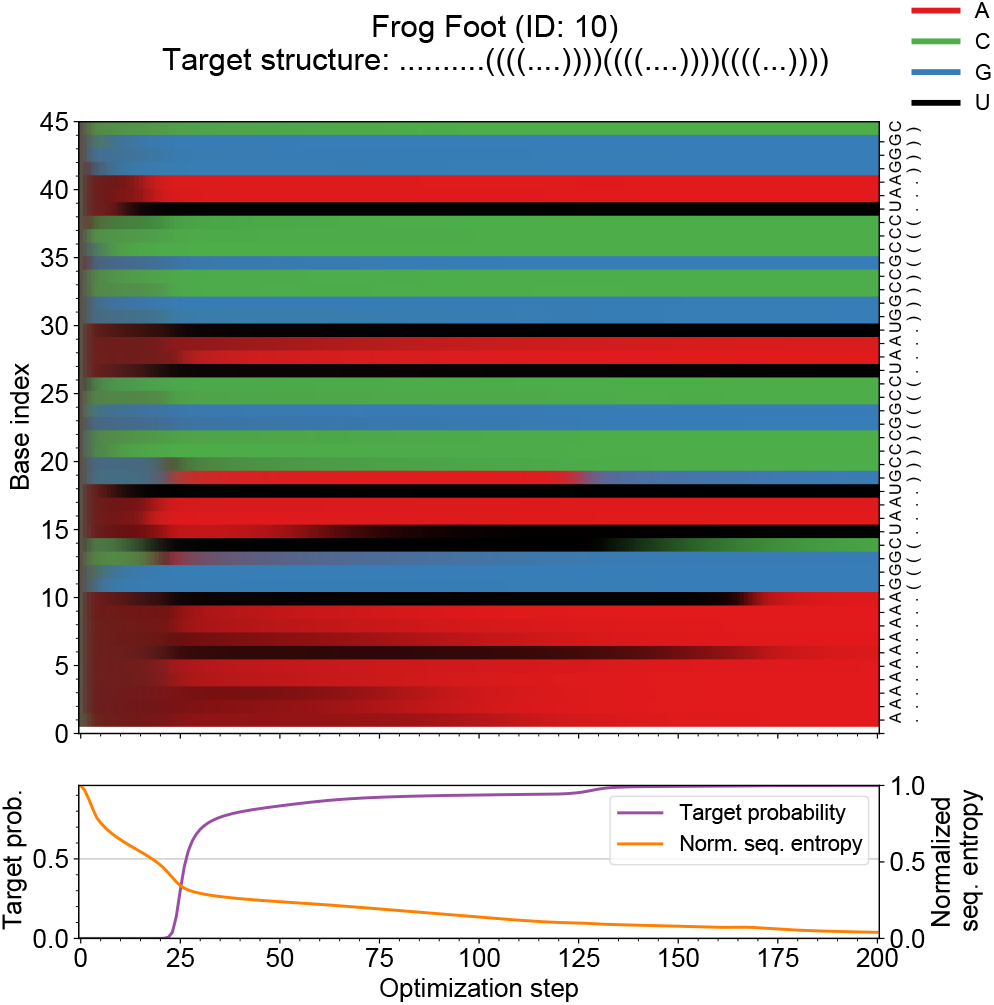
Overview of sequence optimization run for puzzle 10 (“Frog Foot”). Evolution of sequence probability distributions (above), probability of target structure and normalized sequence entropy (below) over the course of an optimization run. The sequence probability distributions at each site are projected into RGB-space and represented as colors. On the right hand side of the sequence probability distributions is the sequence of most likely bases as well as the minimum free energy (MFE) structure predicted for this sequence. This figure shows a successful optimization run with a final target probability of 0.962, with base identity changes happening throughout the optimization run.

In optimization runs of intermediate or low success as measured by final target probability, as shown in Figures 3 and 4 for puzzles 47 and 41, we still see improvements happening over the course of the optimization run, albeit from a low base of target probability, indicating that the optimization run might be stuck in a local minimum.

**Figure 3:**
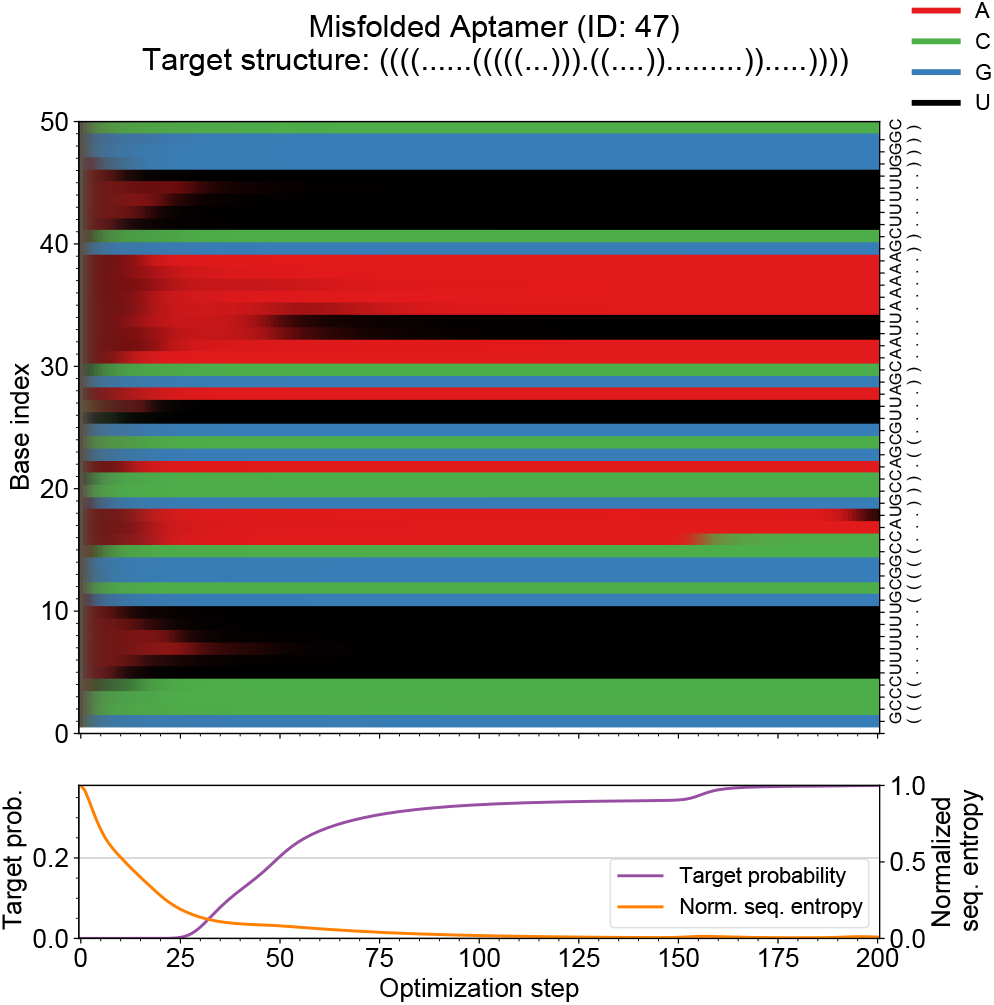
Overview of sequence optimization run for puzzle 47 (“Misfolded Aptamer”). This figure shows an optimization run with a final target probability of 0.381. The designed sequence of most likely bases has the correct minimum free energy structure, but the probability is only 0.381, indicating the possibility for better sequences to exist. Normalized sequence entropy drops off very quickly in the first 50 steps, and between steps 50–150 seemingly little happens. During steps 150–200, there are two base transitions in positions unpaired in the target structure, giving a small increase in target probability.

**Figure 4:**
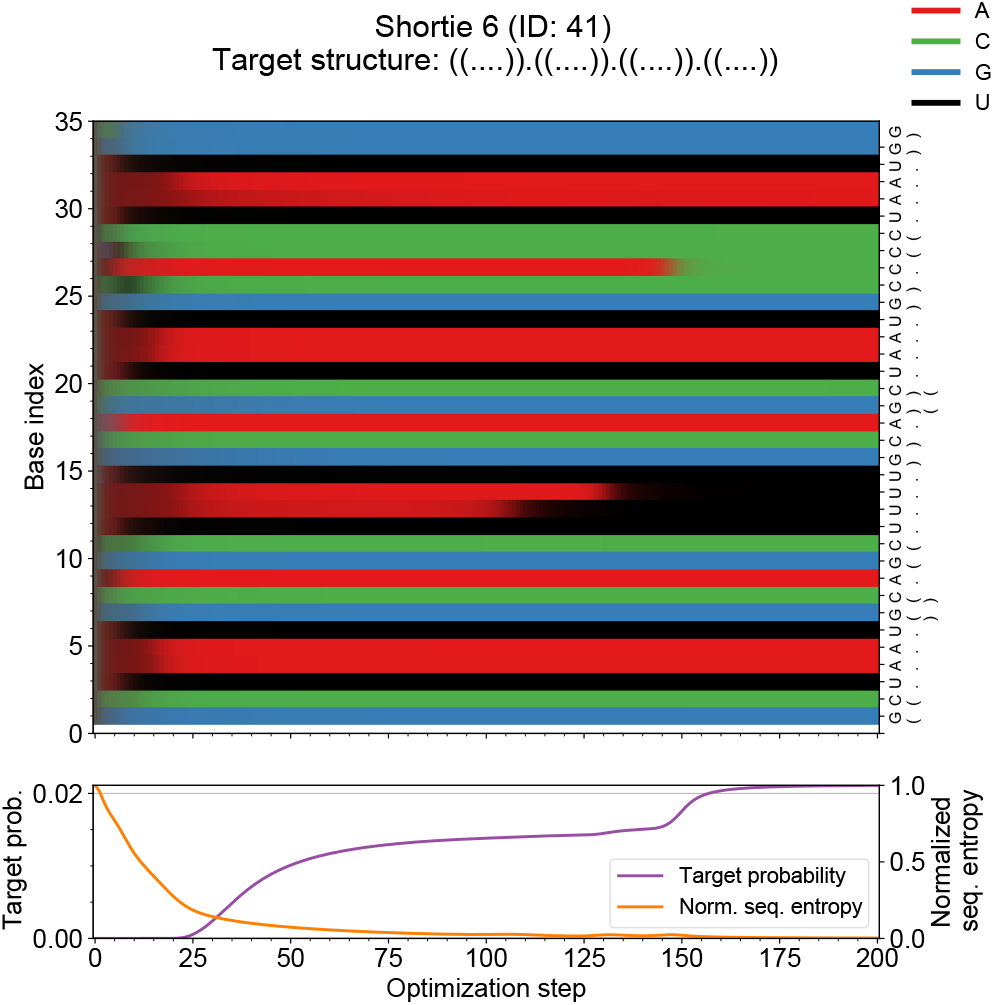
Overview of sequence optimization run for puzzle 41 (“Shortie 6”). This figure shows an optimization run with a final target probability of 0.021. The normalized sequence entropy drops quickly during the first 30 steps, fixing the base identity at all positions that are base-paired in the target structure. The A to C transition at position 27 around step 150 gives a small improvement to target probability.

The results show that our method was successful in computing meaningful gradients. In addition, straightforward gradient optimization was surprisingly effective for the inverse folding problem.

## 4 Discussion

The goal of this work was to develop a differentiable means of computing the partition function for an RNA sequence. To this end, we introduced a new differentiable generalization of McCaskill’s algorithm that accepts probabilistic sequences as input (see Section 2.2). Ideally, our algorithm should be practically useful as well as theoretically sound. There is no theoretical guarantee that the gradients computed via this procedure will be useful for numerical optimization. The results presented in Section 3 provide empirical evidence for this. While there are many relevant loss functions of the partition function over which one could optimize, inverse folding is a straightforward and practical demonstration of our method. Moreover, the arrival at local minima via our simple optimization procedure is an expected property of gradient descent and suggests that combining our method with additional design strategies (e.g., an adaptive walk, multiple initializations) could lead to a breakthrough in this domain.

Given the gradients of the structure-sequence partition function with respect to sequence probability, it is not immediately apparent that gradient descent should always collapse the sequence probability distributions to a one-hot encoded sequence, yet this is what we mostly observed in our optimization runs. While this is certainly a desirable property, it could be advantageous to stop an optimization run earlier in order to sample from the resulting sequence distribution, thereby increasing the number of candidate solutions one can get from one run.

The rapid drop of normalized sequence entropy during the initial stages of optimization corresponds to a fixing of base identities at many positions that are base-paired in the target structure. These positions often do not change their base identity throughout an optimization run and bias the further search to a local minimum in sequence space. This could be improved by performing smaller gradient updates in the initial phase of sequence optimization, before the choice for a specific minimum has been made.

## 5 Future Work and Limitations

The primary limitation of this work is the memory cost of the current implementation. As discussed in Section 2.2, there are many subtle design choices in the recursions (e.g. the treatment of dangling ends and mismatches). In future work, we plan on implementing more memory-efficient variants of these recursions to scale our method to longer sequences. We expect this will be possible given that the approximation for dangling ends, terminal mismatches, and coaxial stacking used by ViennaRNA leads to poor memory usage when adapted to probabilistic sequences. By using a non-approximate model, we could actually use less memory.

Given a means to calculate gradients for sequence lengths of practical importance, there are many applications of our method for the inverse folding of RNA sequences with desired thermodynamic properties. For example, extensions of our method for designing sequences with desired secondary structure could be used for the design of mRNA molecules, such as those used in mRNA vaccines, with specified sequence and structure constraints [27]. Additionally, since we optimize over probabilities, our method could also be used to optimize for more complex equilibrium properties such as bistability. It is also straightforward to optimize for thermodynamic properties other than probabilities of equilibrium structures; for example, one could modify the procedure described in Section 2.3 to optimize for sequences with a desired heat capacity.

Instead of using a fixed energy model and optimizing over the sequence to minimize some function of the partition function, one could equally fix a set of sequences and optimize over the parameters of the energy model to maximize agreement with experimental data [19]. One straightforward application would be a transparent and reproducible reparameterization of the nearest-neighbor model. Equivalently, one could efficiently fine-tune the nearest-neighbor model to chemically modified RNAs, *in vivo* data, or a particular RNA family.

Lastly, we envision our method serving as a loss function in deep learning. For example, Petti *et al*.’s differentiable version of the Smith-Waterman algorithm enabled jointly learning a multiple sequence alignment (MSA) and down-stream parameters (e.g. AlphaFold weights) rather than treating MSA generation as a preprocessing step. Analogously, existing deep learning approaches to RNA structure prediction often use secondary structure as a preprocessing step and could be made end-to-end differentiable via our method [28, 29].

We note that any application of our method to RNA can also be applied to DNA using the Santa-Lucia parameters for the nearest-neighbor model [30].

## 6 Conclusion

The partition function is the fundamental quantity in statistical mechanics and captures the statistical properties of a system in thermodynamic equilibrium. McCaskill’s dynamic programming algorithm for calculating the structure partition function of an RNA sequence allows one to study such properties for a given sequence computationally. However, the reverse is not true; it is not straightforward to use McCaskill’s algorithm to design sequences with desired thermodynamic properties.

In this work, we modify McCaskill’s algorithm so that it is differentiable. This allows one to apply gradient descent over a probability distribution of sequences to optimize with respect to macroscopic thermodynamic quantities. We demonstrate this approach empirically by optimizing for sequences that maximize the probability of folding into a given secondary structure (see Section 3), but note that this method can be adapted to many different optimization problems (see Section 5).

This work serves as an example of the synergies between existing analytical models of physical systems and differentiable programming. The adaptation of existing models to be well-defined over continuous inputs not only serves as a powerful path towards inverse folding, but one in which the relevant quantities are expressive and readily interpretable.

## Software Availability

We have made two implementations of our method publicly available. First, an implementation in JAX using reverse-mode automatic differentiation: https://github.com/rkruegs123/jax-rnafold. Second, an implementation in Julia using forward-mode automatic differentiation: https://github.com/marcom/DiffFoldRNA.jl/.

